# Reversible Polymorphism-Aware Phylogenetic Models and their Application to Tree Inference

**DOI:** 10.1101/048496

**Authors:** Dominik Schrempf, Bui Quang Minh, Nicola De Maio, Arndt von Haeseler, Carolin Kosiol

## Abstract

We present a reversible Polymorphism-Aware Phylogenetic Model (revPoMo) for species tree estimation from genome-wide data. revPoMo enables the reconstruction of large scale species trees for many within-species samples. It expands the alphabet of DNA substitution models to include polymorphic states, thereby, naturally accounting for incomplete lineage sorting. We implemented revPoMo in the maximum likelihood software IQ-TREE. A simulation study and an application to great apes data show that the runtimes of our approach and standard substitution models are comparable but that revPoMo has much better accuracy in estimating trees, divergence times and mutation rates. The advantage of revPoMo is that an increase of sample size per species improves estimations but does not increase runtime. Therefore, revPoMo is a valuable tool with several applications, from speciation dating to species tree reconstruction.

## 1. Introduction

Molecular phylogenetics seeks to understand evolutionary phenomena such as speciation dynamics and biodiversity by estimating evolutionary parameters at the species level. The reconstruction of the species history gives insights into the basic mechanisms of biology. However, the topology of the species tree is not always clear, especially when phylogenies from different genomic regions (i.e., gene trees or genealogies) differ from each other (Degnan and Rosenberg, 2006).

Statistical approaches to tree reconstruction such as maximum likelihood and Bayesian methods rely on substitution models (Tavaré, 1986). These models describe and quantify the probabilities of how sequences may evolve along a phylogeny. They are defined by an instantaneous rate matrix ***Q*** that contains the substitutions rates between the different character states. For computational convenience, most substitution models are *reversible*. That is, the process describing the evolution of the sequence is independent of the direction in time. Reversibility is important in phylogenetics for tree inference from large data sets with many species because it simplifies the likelihood function (Yang, 2006, p.34) and reduces the number of trees by a factor of 2*l* − 3, where *l* is the number of tips of the tree (Hein et al., 2004, p. 70). Finally, rate matrices of reversible substitution models have real eigenvalues (Kelly, 1979) which enables a fast and stable eigendecomposition during matrix exponentiation (Golub and Loan, 1996). Many software packages use reversible substitution models (e.g., HyPhy, Pond et al. 2005; PhyML, Guindon et al. 2010 and MrBayes, Ronquist et al. 2012). RAxML (Stamatakis, 2014) and IQ-TREE (Nguyen et al., 2015) additionally offer efficient tree search algorithms for very large phylogenies.

Substitution models, when naively applied to species trees (concatenation methods, e.g., Gadagkar et al., 2005), assume the species or population to be fixed for a specific character state and do not account for effects on the population genetics level such as Incomplete Lineage Sorting (ILS; Maddison, 1997; Knowles, 2009). Incompletely sorted lineages coalesce deep in the tree and their coalescent events do not match the speciation events. The probability of ILS is large and consequently tree reconstruction is difficult if the time between speciation events is short or if the effective population size is large (Pamilo and Nei, 1988). The multispecies coalescent model can be used to quantify the phylogenetic distortion due to ILS. It simulates a coalescent process (Kingman, 1982) on each branch of the species tree and combines these separate processes when branches join together. This model predicts that for specific evolutionary histories the gene trees with highest abundance conflict the species tree topology (anomaly zone; Degnan and Rosenberg, 2009; Degnan, 2013). These are extreme cases where common tree inference methods not accounting for ILS such as concatenation (Gadagkar et al., 2005) or democratic vote (Pamilo and Nei, 1988) fail because they are statistically inconsistent (e.g., Ewing et al., 2008). However, ILS considerably deteriorates estimates already when species trees are not in the anomaly zone (Pollard et al., 2006).

We have recently developed an approach called Polymorphism-Aware Phylogenetic Model (PoMo, De Maio et al., 2013). PoMo builds on top of substitution models but makes use of within-species data and considers present and ancestral polymorphisms thereby accounting for ILS. Similar to multispecies coalescent models it uses multiple sequence alignments of up to several hundred species while allowing for many within-species sequences to infer base composition and mutational parameters. Recently, we applied PoMo to infer species trees (De Maio, Schrempf, and Kosiol, 2015). We showed in a large scale simulation study with various demographic scenarios and evaluation against other state-of-the-art methods like BEST (Liu, 2008), *BEAST (Heled and Drummond, 2010), SNAPP (Bryant et al., 2012) and STEM (Kubatko et al., 2009) that PoMo is approximately as fast as standard DNA substitution models while being more accurate in terms of the branch score distance (Section 3.1). Furthermore, application to great apes data leads to phylogenies consistent with previous literature and also with the geographic distribution of the populations.

Here, we prove the reversibility of PoMo when an associated reversible mutation model (Section 2.3) is used and derive the corresponding stationary distribution. This will open the PoMo approach to a new area of applications because a reversible model can take advantage of existing algorithms that efficiently reconcile the species tree. We will discuss the reversible solution of PoMo, provide connections to the diffusion equation and introduce an implementation in IQ-TREE (Nguyen et al., 2015). Finally, we present a simulation study and an application to real data to demonstrate the performance of the reversible PoMo (revPoMo) and to confirm its relevancy in medium to large-scale tree search.

## 2. Materials and Methods

### 2.1. *DNA Substitution Models*

DNA substitution models assume that a DNA sequence evolves as a series of independent substitution events which replace a nucleotide by another one. Substitutions are modeled as a time-continuous, time-homogeneous Markov process (Yang, 1994). Additionally, the different sites of a sequence are assumed to evolve independently. The four nucleotides *A*, *C*, *G* and *T* form the alphabet 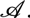. The rates of change *q_xy_* from nucleotide *x* to nucleotide *y* are summarized in an instantaneous rate matrix 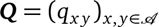 which completely describes the time-continuous Markov process. The assumption of time-homogeneity implies that the entries of *Q* are constant in time. One also assumes stationarity, i.e., the existence of a stationary distribution 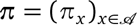 which is the solution to π*Q* = 0. If the Markov process is reversible, then detailed balance *π_x_q_xy_* = *π_y_q_yx_* is fulfilled. Thus, for the General Time Reversible (GTR, Tavaré, 1984) model the rate matrix has the following structure

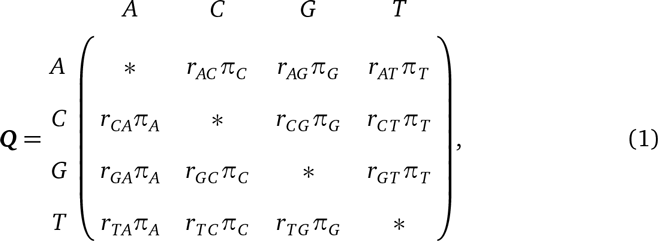

with *q_xy_* = *π_y_r_xy_* and exchangeabilities *r_xy_* = *r_yx_* > 0. The diagonal entries are chosen such that the row sums are zero. The expected number of events on a branch of length *d* is 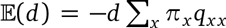. Usually, ***Q*** is normalized such that Σ*_x_*Σ*_x≠y_* π*_x_q_xy_* = 1 or 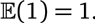

### 2.2. *The Alphabet of revPoMo*

Standard DNA substitution models are limited in the sense that they assume that species are always fixed for a specific nucleotide (i.e., the changes are substitutions). For revPoMo, we use standard DNA models such as HKY (Hasegawa et al., 1985) or GTR (Tavaré, 1984) as mutation models introducing variation into populations that are no longer assumed to be fixed for one nucleotide. We expand the alphabet to include characters that represent polymorphisms so that populations can have polymorphic states. Thereby, revPoMo introduces a virtual haploid population of constant size *N* and distinguishes between fixed (*boundary* {*Nx*} = {*Nx*,*0y*} = {*0y*,*Nx*} and polymorphic characters {*ix*, (*N* − *i*)*y*} (1 ;≤ *i* ≤ *N* − 1; *x*,*y* ∈ {*A*, *C*, *G*, *T*}; *x* ≠ *y*), where *x* and *y* are the nucleotides of the associated mutation model (Fig. 1). For convenience, we call the set of boundary characters the *boundary*. To keep the alphabet of revPoMo 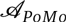 manageable, we assume that at most two different nucleotides per site are present simultaneously. This is only a mild restriction and many real data sets meet this assumption. For example, no sites with three or four nucleotides have been found in the great apes data set described in Section 2.11. This restriction also agrees with the chosen mutation model (Section 2.3). The alphabet-size of revPoMo is

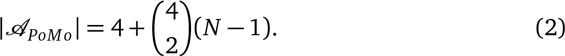

To differentiate between revPoMo and the associated mutation model, we refer to the characters of the mutation model as *nucleotides* and to the characters of revPoMo as *states*. The instantaneous rate matrix of revPoMo *Q_revPoMo_* is composed of the rates of mutations and genetic drift

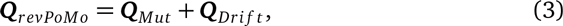

which will be discussed in the next two sections.

### 2.3. *The Mutation Model of revPoMo*

For any nucleotide pair (*x*, *y*) with *x* ≠ *y*, the state {*Nx*} can mutate to state {(*N* − 1)*x*, 1*y*} at rate *μ_xy_* introducing a new nucleotide *y* into the population.

**Figure 1.**
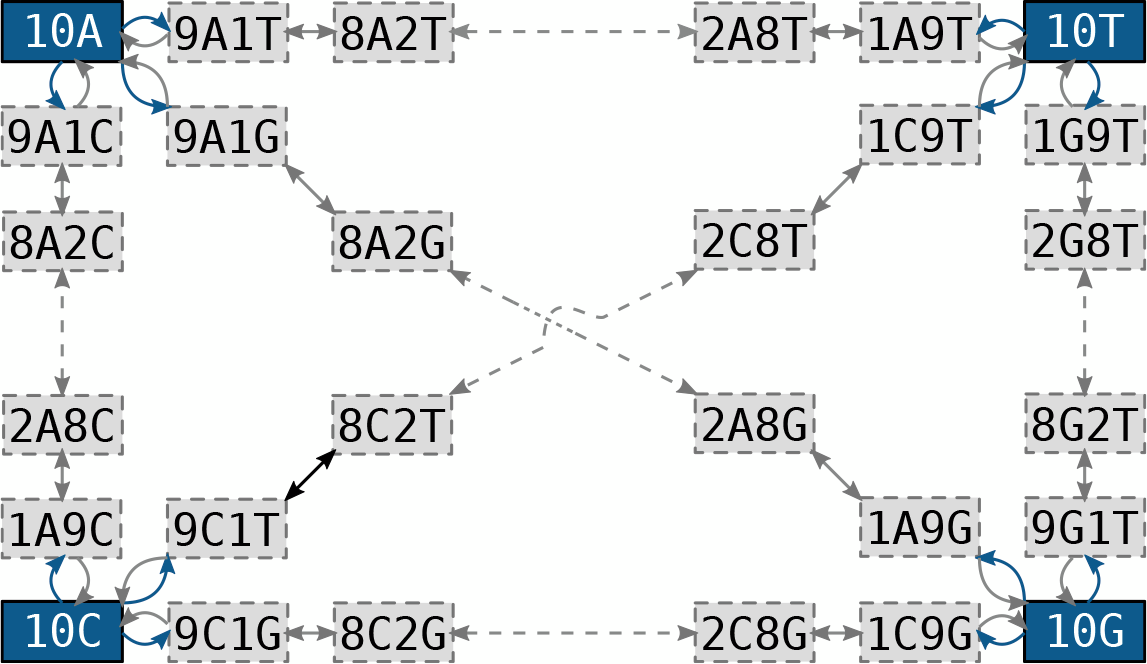
The alphabet of revPoMo and its connectivity for *N* = 10. Blue and gray arrows indicate mutations and genetic drift, respectively. Dashed arrows symbolize the presence of intermediate states. A virtual population that is in the boundary state {10*A*} can move to the polymorphic states {9*A*, 1*C*}, {9*A*, 1*G*} and {9*A*, 1*T*} through a mutational event. Only states with frequency changes of size one are directly connected. For example, two jumps of the Markov process are needed to move from {10*A*} to {8*A*,2*G*}.

Additionally to restricting the state space, mutations are confined to the boundary only. This is a good assumption if mutation rates are low or if genetic drift removes variation reasonably fast (Vogl and Clemente, 2012), a requirement that is met for low effective population sizes. Analogous to the GTR model, the mutation coefficients *μ_xy_* can be decomposed into *μ_xy_* = *m_xy_π_y_*, where *m_xy_* = *m_yx_* and *π_y_* is the entry of the stationary distribution of the mutation model corresponding to nucleotide *y*. Although the concepts are similar we separate the substitution rates from the mutation rates of the associated mutation model by using different symbols (*q_xy_* ~ *μ_xy_*, *r_xy_* ~ *m_xy_*). The symmetry of the coefficients *m_xy_* is a requirement for the reversibility of the GTR model and consequently also of revPoMo. This type of mutation model also fits the structure of the alphabet of revPoMo which only allows two nucleotides to be present in a virtual population. The mutation rates *μ_xy_* of the associated mutation model are summarized in the rate matrix *Q_Mut_* of dimension 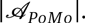. All other rates are zero and the diagonal elements are defined such that the respective row sum is zero.

### 2.4. *Genetic Drift in revPoMo*

The drift rate for a polymorphic state is modeled with the time-continuous neutral Moran model (e.g., Durrett, 2008, p. 46). Given a virtual population of size *N*, in each generation an individual is randomly chosen to reproduce. The offspring is of the same type as the parent and replaces another randomly chosen individual from the population. Thereby, the population size remains constant. For 1 ≤ *i* ≤ *N* − 1, the rate of change from a state {*ix*, (*N* − *i*)*y*} to states {(*i* + 1)*x*, (*N* − *i* − 1)*y*} or {(*i* − 1)*x*, (*N* − *i* + 1)*y*} is

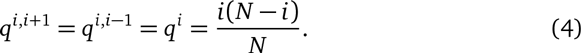

Similar to the mutation model, these rates are summarized in the rate matrix *Q_Drift_* of dimension 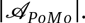. Again, all other rates are zero and the diagonal elements are determined by the requirement that all row sums are zero. For our polymorphic states, the model is symmetric because the rates of increase and decrease are equal. Importantly, nucleotide frequency shifts larger than one require more than one drift event (Fig. 1). In contrast to DNA substitution models, a substitution in revPoMo is the interplay of a mutational event with subsequent frequency shifts such that the newly introduced nucleotide becomes fixed.

### 2.5. *Reversibility of revPoMo*

If the equilibrium of a Markov process exists it is described by the stationary distribution *π* (see above). The stationary distribution of the time-continuous Markov process defined by the instantaneous rate matrix *Q_revPoMo_* (Appendix A) will be denoted *p* to differentiate it from the stationary distribution of the Markov process of the associated mutation model *π*. The entries corresponding to the four boundary states are denoted *p_x_*, the entries corresponding to polymorphic states {*ix*, (*N*−*i*)*y*} (1 ≤ *i* ≤ *N* − 1, *x* ≠ *y*) are 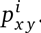. The number of elements of *p* is 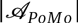 (Eq. 2).

#### Theorem 1.

*The Markov process defined by Q_revPoMo_ is reversible with stationary distribution*

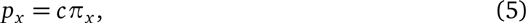

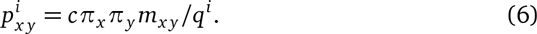

*The normalization constant is*

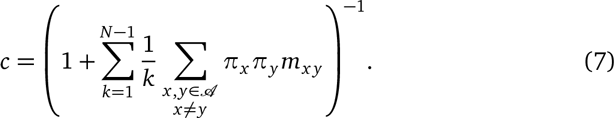

Note that 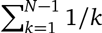 is half of the expected branch length of a genealogy with *N* samples for the standard coalescent model. This coincides with the rate of coalescence in the Moran model being twice as much as the rate in the standard coalescence model. Furthermore, 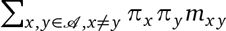 is the expected number of mutations per site and unit time for the associated mutation model.

#### Proof

An irreducible Markov process with a finite number of states is reversible if its (unique) stationary distribution fulfills detailed balance 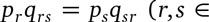 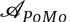; e.g., Norris 1998, p. 125). We distinguish two cases; (a) balance between boundary states and their neighbors

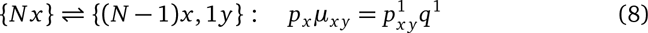

and (b) balance between neighboring polymorphic states

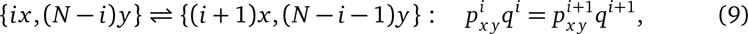

where 1 ≤ *i* ≤ *N* − 2. Both conditions can be verified using Eq. (5) and Eq. (6). Supplemental Section S1 shows the derivation of the normalization constant and the computation of the stationary distribution *p*.

### 2.6. *The Stationary Distribution*

In this section we illustrate the stationary distribution *p* of revPoMo and connect it to previous results of phylogenetics and population genetics. Similarly to DNA substitution models, the frequencies of the boundary states *p_x_* are proportional to the stationary distribution of the nucleotides *π*. revPoMo can estimate this nucleotide distribution empirically from the alignment data or by maximum likelihood. We were concerned that the genetic variation at stationarity of the small, virtual population of revPoMo represented by 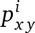 differs considerably from the genetic variation in the real (modeled) populations with high effective population sizes. A substantial amount of theoretical work exists that models the dynamics and the equilibrium properties of populations assuming very large effective population sizes (diffusion limit; e.g., Wright, 1931, Wright, 1945; Kimura, 1964). The diffusion limit has been found to be an adequate approximation in a broad range of population genetics scenarios (Kimura, 1964). We compared the polymorphic entries of the stationary distribution of revPoMo 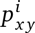 to the stationary solution of the diallelic diffusion equation with drift, equal mutation rates and without selection (e.g., Durrett, 2008, p. 254) which is the probability density of the Beta distribution Ф{*v*, *θ*) ~ Beta{*θ*, *θ*), where *v* is the continuous allele frequency. It converges to 1/*v*(1 − *v*) if the scaled mutation rate *θ* = 4*N_e_μ* is small. The elements of the stationary distribution of PoMo corresponding to polymorphic states 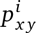 are distributed proportional to its discrete, empirical version 1/*i*(*N*−*i*) (Fig. 2). In particular, revPoMo is a good approximation if *θ* < 0.1. Many real data sets (e.g., genomic sequences of mammals, and *Drosophila* species) meet this requirement. For microbial data sets, however, this assumption might not be valid. Section S2 includes additional thoughts on the stationary distribution for non-uniform *π*.

**Figure 2.**
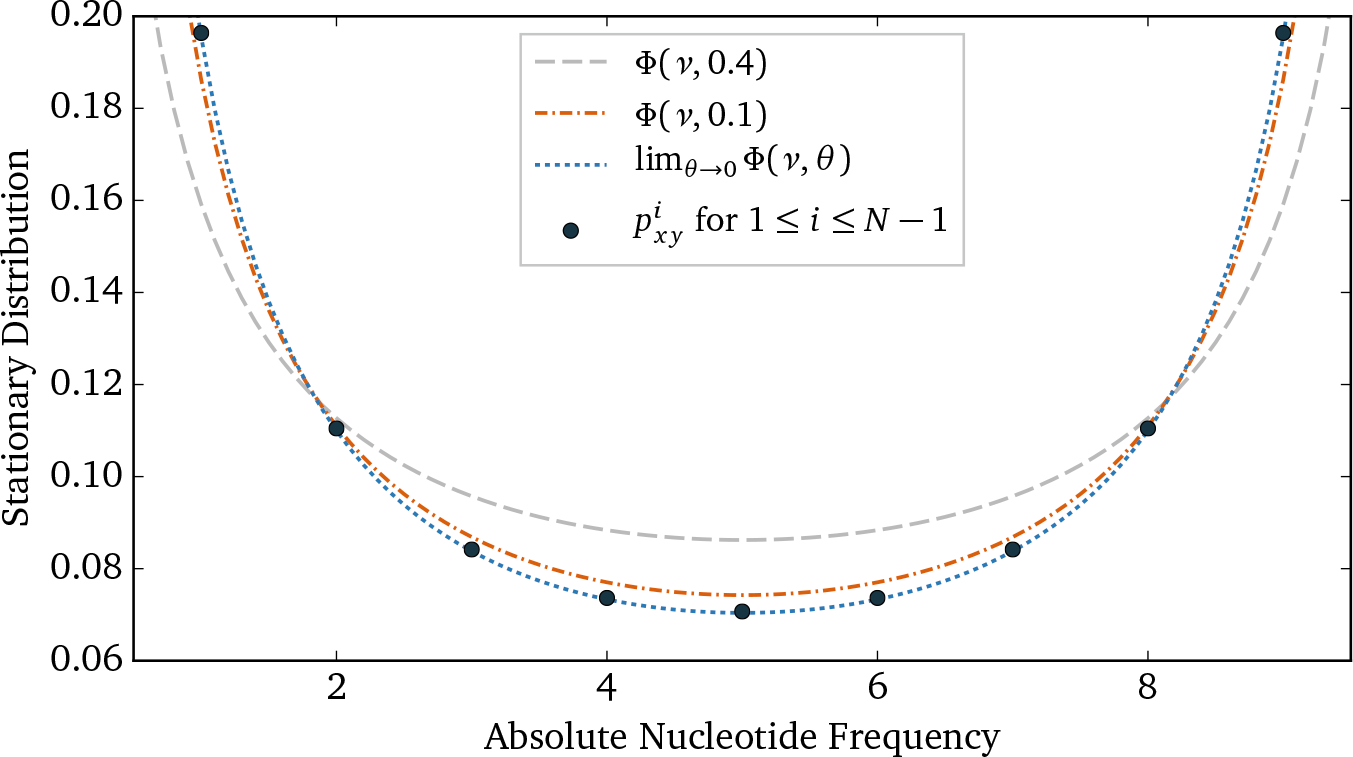
A comparison of the stationary solution of the diffusion equation (Wright, 1931) Ф{*v*, *θ*) = *v*^θ−1.0^(1 − *v*)^θ−1.0^ for equal scaled mutation rates *θ* = 4*N_e_μ* and drift with the polymorphic elements of the stationary distribution of revPoMo for *N* = 10. *v* is the continuous relative nucleotide frequency. Both *p* and Φ have been normalized such that the polymorphic elements integrate to one and the domain of Φ has been expanded from (0,1) to (0,10). Ф converges to a continuous version of 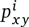 if *θ* goes to zero (blue dotted line) because revPoMo only allows for mutations at the boundaries.

### 2.7. *Number of Parameters*

This section discusses a peculiarity of the number of parameters of revPoMo compared to the associated mutation model. For example, both the GTR model and revPoMo with the GTR model as associated mutation model (GTR + revPoMo) have three free parameters for the stationary nucleotide frequencies because the four entries of *π* have to sum to one. Furthermore, the GTR model normalizes the substitution coefficients such that the total substitution rate is one per unit time, i.e.,

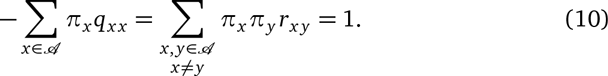

Thereby it reduces the number of free rate parameters from six to five. This step is necessary because only the ratios of the substitution coefficients matter and the total substitution rate between nucleotides is confounded with the branch lengths. However, in revPoMo the total rate of mutations cannot be constrained in the same way because it also determines the percentage of polymorphic states in the stationary distribution *p* (Eq. 7). Scaling the symmetric mutation coefficients *m_xy_* by a common factor affects the ratio 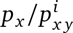. This is why the number of parameters of revPoMo is larger by one, e.g., the GTR model has eight and GTR + revPoMo has nine parameters. Previously, this additional unknown variable was empirically estimated (De Maio et al., 2013). In contrast, we jointly infer it with all other model parameters.

### 2.8. *Relation between revPoMo and Substitution Models*

It is desirable to compare distance estimations of revPoMo with estimations from standard DNA substitution models. We have seen however, that a mutation from the boundary requires subsequent nucleotide frequency shifts to become a substitution and that the total number of mutations scales with phylogenetic distance. Phylogenetic distances are usually normalized such that on average one event (i.e., one jump of the Markov process) happens per unit length (see also Section 2.7). If a Markov process starts in equilibrium, we have 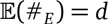, where *$_E_* is the number of events and *d* is the total branch length. For substitution models, the expected number of substitutions is 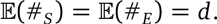. Also in revPoMo the total rate of events per unit length can be normalized to one. However, events can still be either mutations or frequency shifts. Let 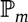 be the probability that an event is a mutation. This mutation moves away from a boundary state {*Nx*} towards a different nucleotide *y*. Let *h* be the hitting probability of the opposite boundary state {*Ny*} before moving back to {*Nx*}. *h* does not depend on *x* and *y* because revPoMo assumes boundary mutations only. The expected number of substitutions is

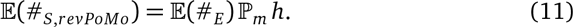

From population genetics, we know that *h* = 1/*N* (e.g. Ewens, 2004, p. 105) because genetic drift is the only active force and the frequency of *y* is just 1/*N*. A comparison of the transition rates of *Q_revPoMo_* shows that also 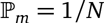 (Appendix B) and we get

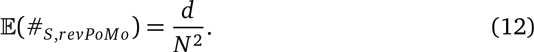

This enables us to compare the branch lengths of revPoMo with the ones of standard DNA substitution models if we assume that the estimated number of substitutions 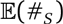 and 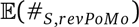 are equal across both models.

### 2.9. *Implementation*

We present an implementation of revPoMo in IQ-TREE (Nguyen et al. 2015; for technical details see Section S5). We allow the virtual population size *N* to vary between 2 and 19. The maximum of 19 is an arbitrarily chosen cut-off to keep the size of the executable small. revPoMo uses multiple sequence alignments in the form of counts files as input data (De Maio et al., 2015). That is, nucleotide counts are given for each site and population. In general, the nucleotide counts (i.e., the number of sampled individuals from a population) will differ from the virtual population size *N* of revPoMo. Furthermore, sequencing errors, merged data from different sources as well as alignment problems may lead to a variation of nucleotide counts between populations or even within populations at different sites. In contrast to DNA substitution models, where the character of the corresponding terminal node is set to the observed nucleotide, the revPoMo state at the same terminal node is not obvious anymore if the sample size is not equal to *N*. A simple method to determine the revPoMo state is to sample *N* nucleotides with replacement from the given data. We do this independently for each site and population and call this method *sampled*.

Instead, similarly to handling ambiguity and error (Felsenstein, 2004, p. 255), we can also weight the revPoMo states at each terminal node according to their likelihood of representing the observed counts. We set these likelihoods to the binomial distribution because a revPoMo state represents a real population with the same proportions of nucleotides. In detail, for a terminal node with observed nucleotide counts {*jx*, (*M* − *j*)*y*}, the likelihood of revPoMo state {*ix*, (*N* − *i*)*y*} is

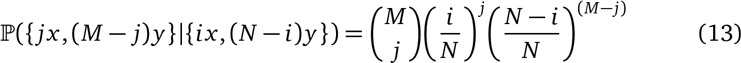

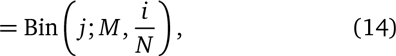

where 0 ≤ *j* ≤ *M* and 0 ≤ *i* ≤ *N*. Only nucleotides that are present in the virtual population can be sampled. We call this sampling method *weighted*.

### 2.10. *Simulation Study*

The performance of revPoMo was tested with simulated sequences. Two different pipelines were used to create genealogies. First, for scenarios with four (e.g., Fig. 3) and eight extant species, the species trees were predefined and genealogies were simulated with the coalescent simulation program MSMS (Ewing and Hermisson, 2010). For the coalescent simulations, the tree height is specified in units of effective population size *N_e_*. We used values of 1 *N_e_* and 10 *N_e_*. Between two and twenty individuals were sampled per population. That is, the genealogies for the scenarios with four taxa have up to 80 tips. Second, in the larger scenario with 60 species, the species trees were created under a Yule birth model (Yule, 1925). A new species tree was simulated for each replicate. We used SimPhy (Mallo et al., 2015) to simulate the species tree and subsequently the genealogies with ten individuals per population. Overall, these genealogies have 600 tips. The tree height was fixed to 3 *N_e_* and the parameter describing the rate of speciations events per coalescent time unit was determined such that the expected coalescence time for 60 species matches 3 *N_e_*. This corresponds to a rate of 1.226 speciations per *N_e_*.

**Figure 3.**
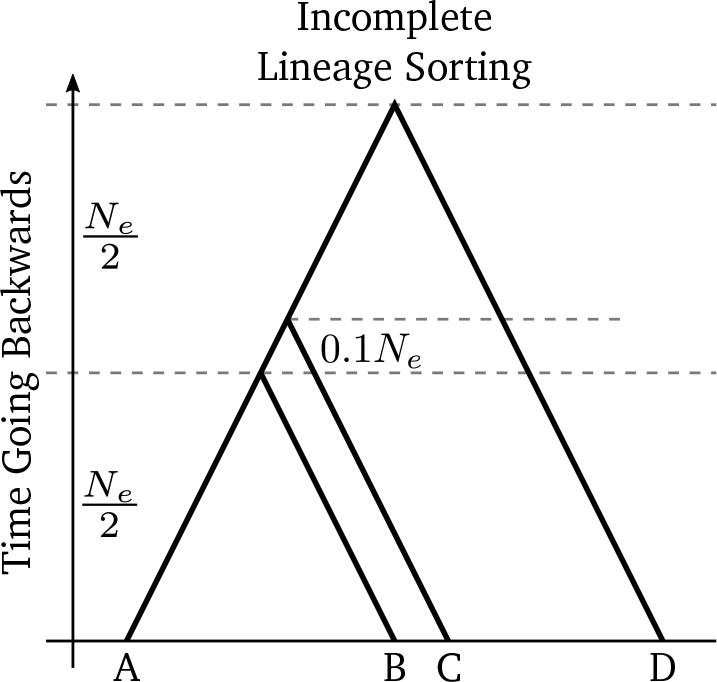
The *Incomplete Lineage Sorting* (ILS) scenario is a species tree with four tips and height 1 *N_e_*, where *N_e_* is the effective population size. The lengths of non-labeled branches are defined by the strict-molecular clock constraints. Using the multispecies coalescent model, we expect about 50 percent of the simulated genealogies to exhibit ILS.

For both pipelines 1000 independent genealogies were simulated. We used these genealogies to generate DNA sequences of genes with Seq-Gen (Rambaut and Grass, 1997) using the HKY model. Thereby, we scaled the branch lengths with a factor of 0.0025. In particular, for scenarios with tree height 1 *N_e_*, the number of substitutions per site from the root to the tips is 0.0025 on average. Likewise, the level of polymorphism within a species was equal for all scenarios and corresponds to an average Watterson’s theta (Watterson, 1975) of 0.0025 per site. The sequence or gene length per genealogy was set to 1000 base pairs (bp). This simulation procedure is equivalent to no recombination within genes, free recombination between genes and no migration between species. The amount of input data was varied between three and all 1000 genes. We performed ten replicate analyses for each setting. In the main text, we include results for the *Incomplete Lineage Sorting* species tree scenario (Fig. 3; De Maio, Schrempf, and Kosiol, 2015) and Yule trees with 60 species. A calculation according to the multispecies coalescent model shows that for the incomplete lineage sorting scenario, if one gene is sampled out of each species *A*, *B* and *C*, about 55% of the genealogies that connect these genes are expected to exhibit ILS.

The analysis of the simulated data focuses on the comparison of three different methods: (1) a standard concatenation approach, where the input sequences within species are concatenated and a DNA substitution model is used for the analysis (2) the non-reversible PoMo with *N* = 10 implemented in HyPhy (Pond et al., 2005; De Maio et al., 2015) and (3) the new reversible version with varying *N* and *weighted* sampling implemented in IQ-TREE (Nguyen et al., 2015). All methods use the HKY substitution model (Hasegawa et al., 1985).

The accuracy of the estimation was measured with the branch score distance (BSD, Kuhner and Felsenstein, 1994) between the true and the estimated species tree. The BSD is the square root of the sum of the quadratic differences in branch lengths between two trees. For one taxon trees it coincides with the relative branch length error. Branches that do not exist in both trees due to differences in topology fully contribute to the BSD. Before we calculated BSDs with PHYLIP (Felsenstein, 2005), the trees were normalized such that their total branch lengths equal 1.0. Normalization is necessary because branch lengths are confounded with substitution rates for DNA substitution models and have a different meaning for PoMo (Section 2.8). Section S3 and S4 provide command lines for the simulation and analysis procedure, respectively.

### 2.11. *Application to Great Apes*

Shared ancestral polymorphisms are very common in great apes (Dutheil et al., 2009). A variety of evolutionary patterns, short internal branches as well as closely related taxa lead to a high level of incompletely sorted lineages between Humans, Chimpanzees and Gorillas (about 25%, Scally et al., 2012). We apply revPoMo to a data set that includes all 6 great apes species divided into 12 populations (Prado-Martinez et al., 2013). The number of sequences per population varies highly between 1 (*Gorilla gorilla diehli*) and 23 (*Gorilla gorilla gorilla*). About 2.8 million exome-widely distributed, 4-fold degenerate sites were analyzed. We use the *sampled* input method which may lead to differences in estimates between runs. We do not expect a high divergence between runs but asses the variance by doing ten replicate analyses.

**Figure 4.**
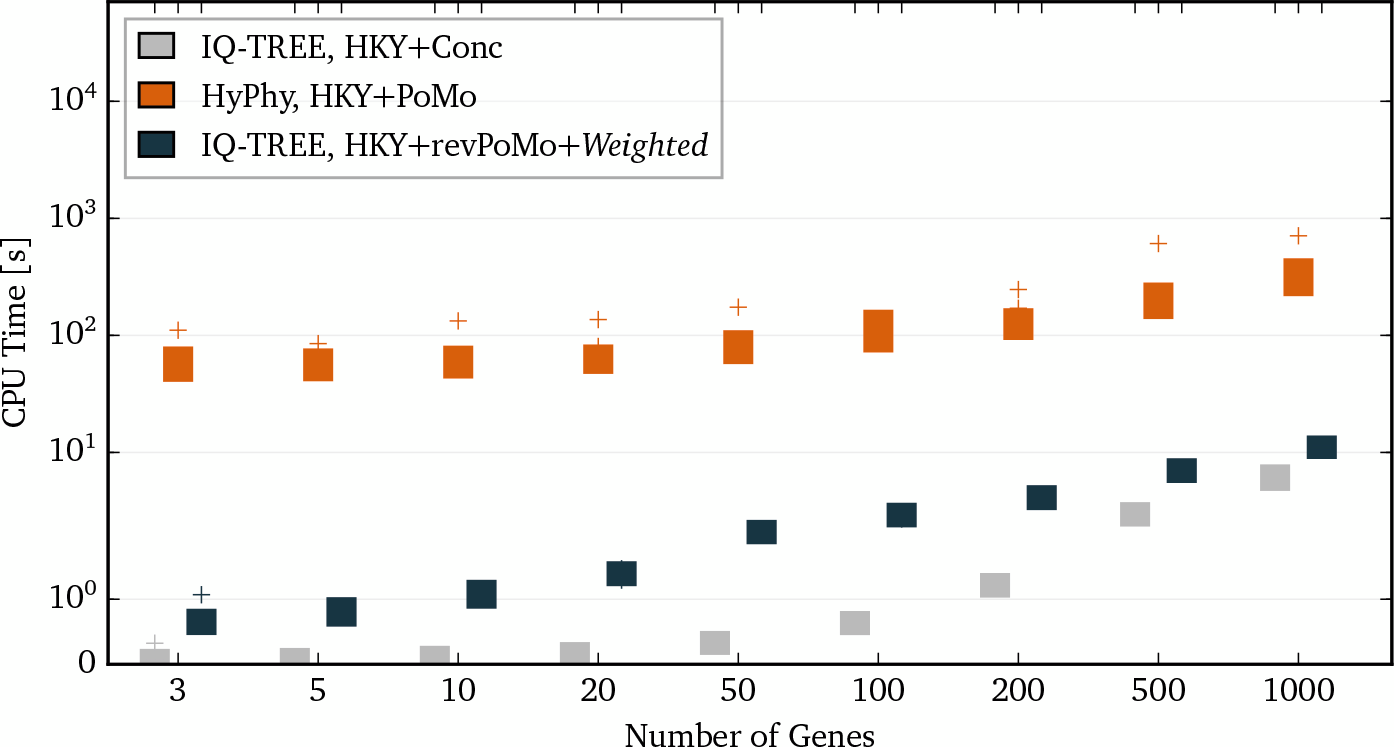
A boxplot of the runtimes of the concatenation approach (IQ-TREE, HKY+Conc), the nonreversible PoMo with *N* = 10 (HyPhy, HKY+PoMo) and revPoMo with *N* = 10 and the *weighted* sampling scheme (IQ-TREE, HKY+revPoMo+*Weighted*) for the ILS scenario with ten samples and a tree height of 1 *N_e_* (Fig. 3). The HKY model was used for all methods. Ten replicate analyses were performed. Different amounts of input data are shown on the x-axis (each gene has a length of 1000 bp).

## 3. Results and Discussion

### 3.1. *Simulations*

A previous simulation study showed that the non-reversible PoMo outperforms other state-of-the-art methods (De Maio, Schrempf, and Kosiol, 2015) in estimating species trees from large data sets. To begin with, we assay the speed of the concatenation method, the non-reversible PoMo and revPoMo for different amounts of sequence data (in number of genes, one gene has 1000 bp). The introduction of reversibility improves speed greatly and the new implementation of revPoMo in IQ-TREE runs up to 50 times faster than the the version implemented in HyPhy. Overall, the runtime is similar to that of standard DNA substitution models (Fig. 4 and Section S6 with Fig. S1-S3).

**Figure 5.**
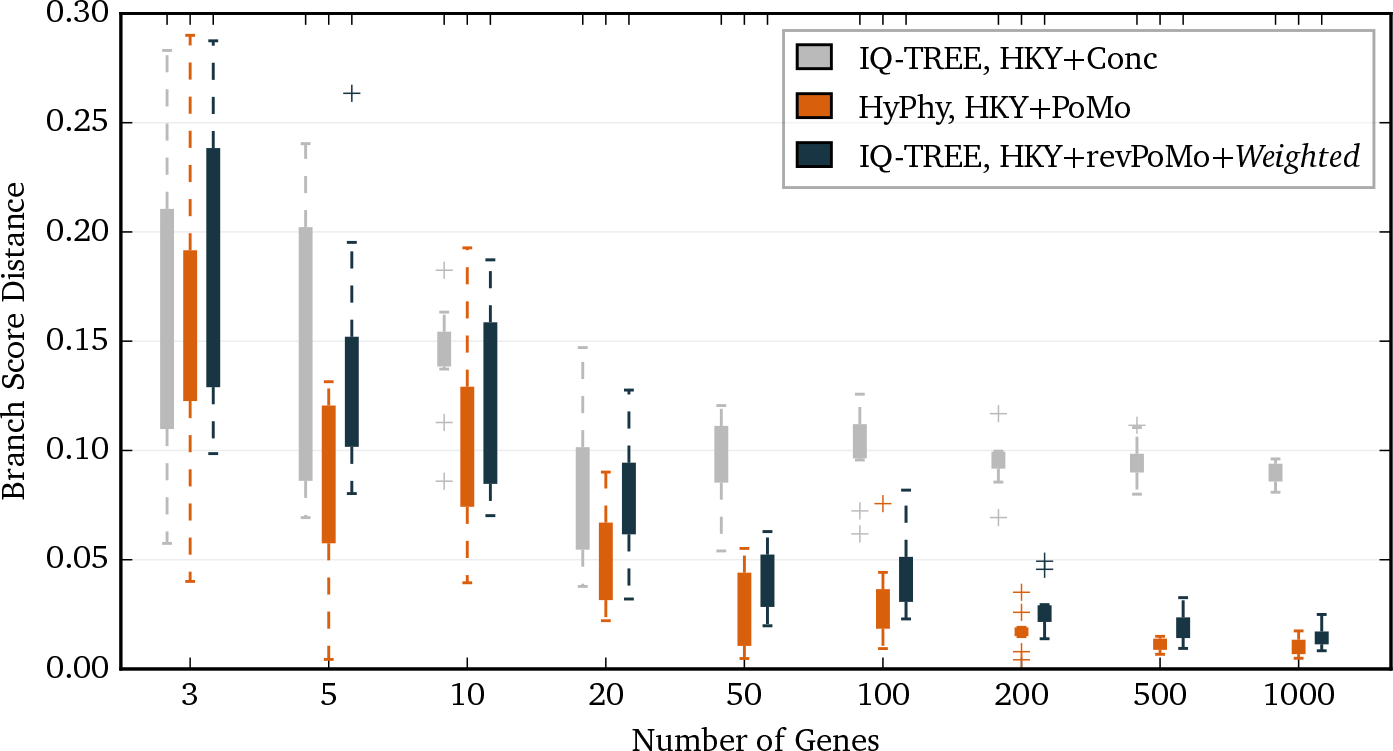
Tree error measured by the branch score distance for concatenation (IQ-TREE, HKY+Conc), the non-reversible PoMo with *N* = 10 (HyPhy, HKY+PoMo) and revPoMo with *N* = 10 and the *weighted* sampling scheme (IQ-TREE, HKY+revPoMo+*Weighted*) in dependence of the amount of data; one gene has 1000 bp. The HKY model was used for all models. The analyzed sequences were simulated under the ILS scenario with ten samples and a tree height of 1 *N_e_* or 0.0025 substitutions per site (Fig. 3). The non-reversible version performs marginally better because the frequency distribution at the root is arbitrary.

The simulation scenario with four species exhibits a significant amount of ILS and both PoMo approaches outperform the concatenation method if the input data contains enough independently evolved genes (Fig. 5). At least 50 genes (50k bp) are needed to get trustworthy results with small standard deviations and analyses of 1000 genes have an error of about 2% only. In general, the error is small (Section S7 and Fig. S4-S18). For species trees that do not exhibit any incomplete lineage sorting, the accuracy of PoMo measured in BSD is similar to the one from concatenation methods and slightly better if more than three samples per population and about 50 genes are available (e.g., Section S7.3 and S7.4 but also De Maio et al., 2015).

With the non-reversible version of PoMo we were limited to trees of about a dozen species only. revPoMo takes advantage of efficient algorithms and the reduced runtime enables us to analyze trees with many species. Here, we present an analysis of trees with 60 species generated under the Yule birth model. The non-reversible PoMo approach is too slow for trees of this size. The runtime of revPoMo on sequences with 1000 genes is about 4.5 h with a standard deviation of about 25min (i5-3330S, 2.70GHz, 2 physical cores). Taking polymorphisms into account improves accuracy in terms of BSD for this scenario. In particular, if more than 100 independently evolved genes are used for the analysis, the BSD is reduced by a factor of seven (Fig. 6). Notably, revPoMo performs better than concatenation methods already if three genes are available. Although it is expected that an increase in the number of species leads to a higher chance of topological errors, the total error is similar to the one of the ILS scenario with four species only. Section S7.5 includes results for a Yule tree with 50 species.

**Figure 6.**
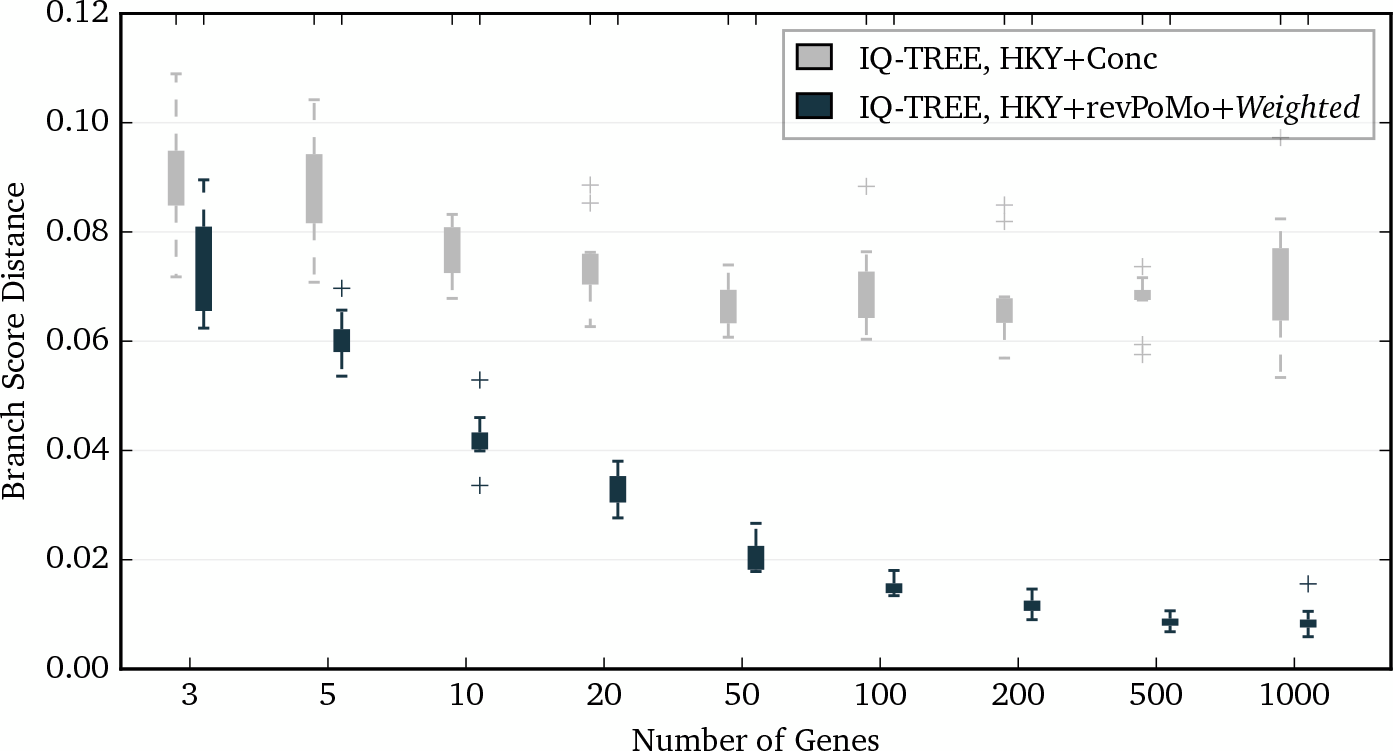
Branch score distance for the concatenation approach (IQ-TREE, HKY+Conc) and revPoMo with *N* = 9 and the *weighted* sampling scheme (IQ-TREE, HKY+revPoMo+*Weighted*) applied to sequences simulated under a Yule tree with 60 species with ten samples each. The HKY model was used in both cases. The tree height is 3 *N_e_* and the level of polymorphism measured by Watterson’s theta is *θ_W_* ≈ 0.0025 per site. Ten replicate analyses were performed. The *x*-axis denotes the number of genes that were analyzed (one gene has 1000 bp).

A very important variable of PoMo is the virtual population size *N* which has initially been set to ten for parameter estimation (De Maio et al., 2013). Up to this point, only the weighted sampling method has been used. Now we use both sampling methods *sampled* and *weighted* to analyze the ILS scenario with ten sequences per species and 1000 genes in dependence of *N*. We find that the allowance of a single polymorphic state for each pair of bases (*N* = 2) already decreases the tree estimation error and that an increase of *N* from two to nine greatly improves the accuracy (Fig. 7). During a further increase of *N* up to 19 the improvement is only marginal. Random sampling with replacement of *N* samples from the data gives better results if *N* is very low. For higher virtual population sizes between 5 and 15, weighting the partial likelihoods performs better on average and is also numerically more stable. Values of *N* above the sample size do not add useful information and therefore do not positively influence the performance.

**Figure 7.**
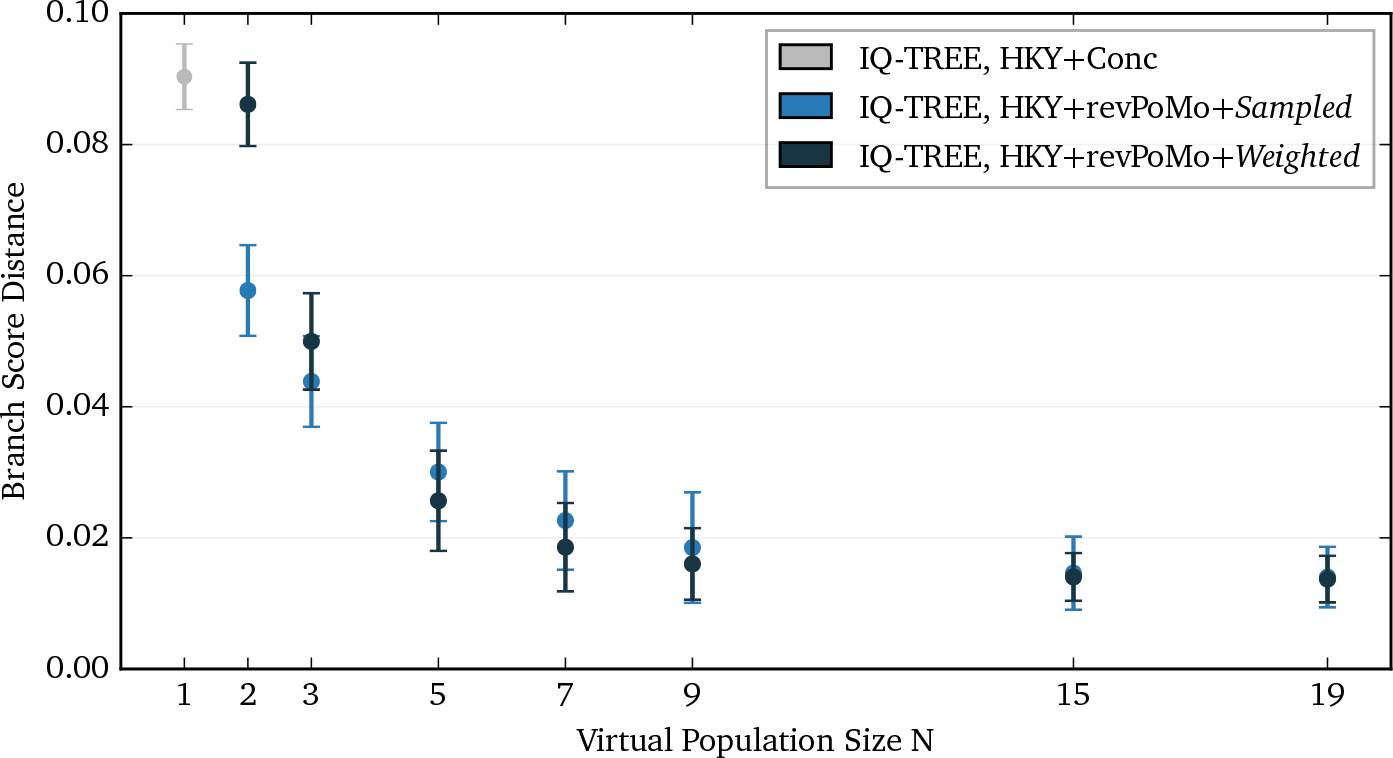
The branch score distance in dependence of the virtual population size *N* for revPoMo with both sampling techniques (IQ-TREE, HKY+revPoMo+*Sampled*; IQ-TREE, HKY+revPoMo+*Weighted*). The analyzed scenario is incomplete lineage sorting with ten samples, a tree height of 1 *N_e_* and 1000 genes of input data. The error bars are standard deviations of ten runs. For *N* = 1, the estimate of the concatenation approach (IQ-TREE, HKY+Conc) is shown. All models use the HKY model.

Additional results (Section S8 and Fig. S20-S24) confirm that for the *weighted* sampling method an increase of the virtual population size above the sample size does not greatly improve the results. For the *sampled* input method and *N* above ten, we also observed numerical underflow errors due to low frequencies of polymorphic states of the stationary distribution if the alphabet is oversized. In general, we advice to choose *N* between five (large trees) and 19 (small to intermediately sized trees), depending on computational resources, tree size and input data. The *sampled* input method seems to do better if the average number of samples is below three.

**Figure 8.**
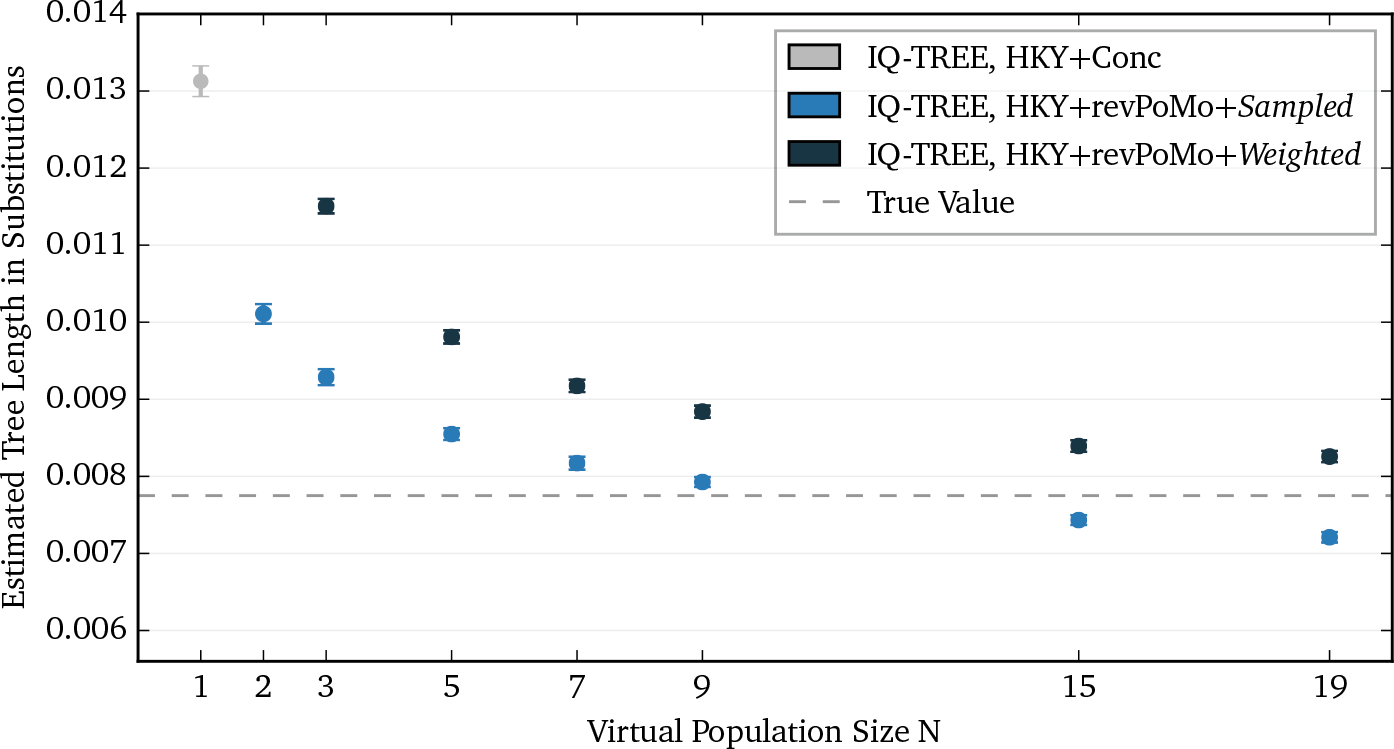
The estimated tree length in substitutions per site in dependence of the virtual population size *N* for revPoMo with both sampling methods (IQ-TREE, HKY+revPoMo+*Sampled*; IQ-TREE, HKY+revPoMo+*Weighted*). The analyzed scenario is incomplete lineage sorting with ten samples, a tree height of 1 *N_e_* and 1000 genes of input data. The errors bars are hardly visible and denote standard deviations of ten replicate analyses. The dashed line is the true value. For *N* = 1, the estimate of the concatenation approach (IQ-TREE, HKY+Conc) is shown.

The total branch length of the inferred phylogeny is a further criterion to judge the quality of revPoMo. Usually, the branch lengths of phylogenies inferred by Markov process based models are given in units of *estimated average number of events per site*. The connection between mutation and substitution rates (cf. Methods) allows an interpretation of the estimated branch lengths of revPoMo. In particular, we can convert the branch lengths to *estimated average number of substitutions per site*, compare them to estimations from standard substitution models and — for simulations — also to the true value (Fig. 8).

The concatenation approach systematically overestimates the phylogenetic distance because polymorphisms are interpreted as substitutions. For revPoMo, we find that the estimated tree length in substitutions improves for both input methods if *N* is increased. The *sampled* input method seems to converge faster but overshoots for values of *N* above the sample size. The further decrease of branch lengths can be attributed to an unnecessary interpretation of substitutions as standing polymorphisms. We conclude that it is only preferable to use the *sampled* input method when the data contains populations with very few individuals.

**Figure 9.**
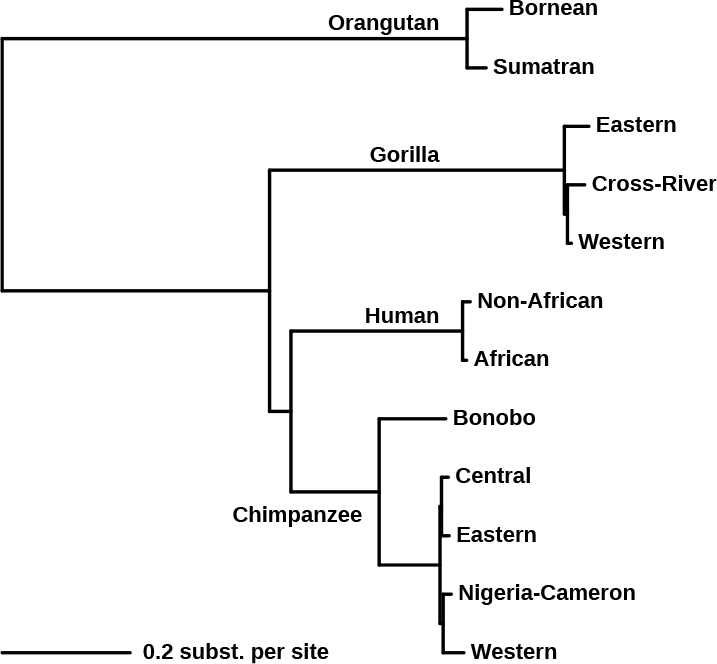
The estimated phylogeny of the great apes data set with revPoMo and the GTR model agrees with the geographic distribution of the species. There are no topological differences between the ten replicate analyses. The virtual population size was set to *N* = 9 and the input method to *sampled*. The phylogenetic scale is in substitutions/site and can be directly compared to values inferred by standard substitution models.

### 3.2. *Real Data*

The previous, non-reversible PoMo already performed well on the great apes data set (De Maio et al., 2015). The phylogeny estimated by revPoMo (Fig. 9) agrees with the geographic distribution of the great apes (species with neighboring habitats are more closely related than species that live further apart) and the topology presented in the original publication (Prado-Martinez et al., 2013). revPoMo evaluates all (*weighted*) or nearly all (*sampled*) available polymorphic information in the data and we expect that estimates between consecutive runs have no or low variance, respectively. Indeed, ten replicate analyses with the GTR model (Tavaré, 1986), *N* = 9 and the *sampled* input method show that it is stable and accurate. The estimated topologies are identical and the total branch lengths have a mean of 3.08 · 10^−2^ substitutions/site with a very low standard deviation of about 6.45 · 10^−7^ substitutions/site. On the contrary, the topology inferred by DNA substitution models is not stable and depends on the individuals chosen to represent the species (De Maio, Schrempf, and Kosiol, 2015).

The branch lengths of revPoMo can be used to estimate the germline mutation rate per generation within the Human-Chimpanzee-Gorilla clade. This is interesting because many discrepancies of estimates have been discussed in the past (Scally and Durbin, 2012). We assume that Humans split from Chimpanzees 7 million years ago (Ségurel et al., 2014) and that the Human-Chimpanzee clade split from the Gorilla clade 10 million years ago (Scally and Durbin, 2012). Furthermore we set the generation time to 25 years. Then, we get an estimate of about 2.65 · 10^−8^ germline mutations per generation per site. This value lies on the lower boundary of other estimates from phylogenies (Li and Tanimura, 1987; Takahata and Satta, 1997). We stress that this is a rough estimate that ignores various complex aspects considered by Li and Takahata. However, with our approach we take into account the effect of standing and ancestral variation on the estimate of mutation rates.

## 4 Conclusions

Polymorphism-aware phylogenetic models have been shown to improve accuracy substantially in parameter estimation (De Maio et al., 2013) and tree inference (De Maio, Schrempf, and Kosiol, 2015) in the presence of ILS. However, the number of populations that could be analyzed with the non-reversible PoMo implementation was limited. Here, we present a reversible PoMo under the following assumptions: (a) polymorphic states can only contain two different nucleotides, (b) the associated mutation model is reversible, (c) drift is described by the continuous-time Moran model (e.g., Durrett, 2008, p. 46) and (d) mutations can only happen when a nucleotide is fixed in the population. The stationary distribution for polymorphic states mimics the stationary solution of the diffusion equation without selection and low scaled mutation rate *θ*.

The number of free parameters of revPoMo is determined by the associated mutation model plus one for the total mutation rate which determines the proportion of polymorphic states. This additional parameter can also be empirically estimated from the data. A generalization of the mutation model such that mutations can happen anytime not only naturally demands a further expansion of the alphabet of revPoMo to allow states with multiple nucleotides but also introduces problems with respect to reversibility. Because of the Kolmogorov criterion (Kelly, 1979, p. 21), the mutation coefficients themselves have to be symmetric then, i.e., *q_xy_* = *q_yx_* and not only *r_xy_* = *r_yx_* in Eq. (1). This is incompatible with mutation models that use estimates of nucleotide frequencies like the HKY model. Furthermore, for polymorphic states the stationary distribution is symmetric with respect to an interchange of nucleotides (Section 2.4). It may be interesting to investigate if and only if this is symmetry is implied by a reversible mutation model.

The introduction of reversibility slightly increases the error in tree inference for some scenarios that have been examined but greatly improves runtimes up to a factor of 50. This allows the reconstruction of large-scale phylogenies. As an example, a Yule tree with 60 species was analyzed and low error rates were observed. We confirm that revPoMo does well on real data and infers a phylogeny that agrees with the geographic distribution of the analyzed populations. We also presented how the branch lengths of phylogenies estimated by revPoMo can be interpreted and compared to the ones estimated by standard substitution models. Finally, we show how the introduction of polymorphic states and an increase of the virtual population size *N* improves estimates. We advice to choose N between five (large trees) and 19 (small to intermediately sized trees), depending on computational resources, tree size and input data. We discourage from using revPoMo on sequence data where no population data is available yet.

Describing the evolution of DNA sequences with Markov processes is very fast but restricts the possibilities of revPoMo to include, e.g., a model of gene flow. However, we want to assess robustness against gene flow in the future. An extension that we would like to implement is the inclusion of rate variation, for example with a gamma distribution. Rate variation might not only be modeled between sites but also along the tree, e.g., to account for changes in effective population size. In particular, it is of high interest to relate the virtual population size of revPoMo to the effective population size of real populations. This would allow direct inference of effective population size as well as germline mutation rates.

Two different methods to process the data at the leaves of the phylogeny *sampled* and *weighted* were implemented. We found that both sampling schemes influence accuracy, especially when the sample size is low. With the *weighted* sampling method a beta-binomial distribution could be used to allow pool sequence (Futschik and Schlötterer, 2010) input data with sequencing errors (Appendix S5.2). Furthermore, one could run a diffusion process that connects the data to the leaves to improve the determination of the likelihoods of the revPoMo states. This would also enable us to model population genetic effects with large or even variable *N_e_* relatively close to the present which is the stage where these effects are most important. We also would like to enable automatic bootstrap with IQ-TREE. Importantly, we want to stress that the idea of revPoMo can be used with substitution models of any type including alphabets consisting of amino acids or codons.

revPoMo is peculiar in the sense that it is discrete in frequency but continuous in time. This property makes it a connection between models that are discrete in time and frequency (e.g., Wright-Fisher model with mutations) and the diffusion limit which corresponds to continuity in time and frequency. The advantage of revPoMo compared to multispecies coalescence based models is that an increase of sample size improves tree and parameter estimations but does not increase runtime. We believe that revPoMo is a valuable tool in species tree estimation from population data.

## 5. Acknowledgments

We thank Claus Vogl and Asger Hobolth for discussions and valuable comments about revPoMo. This work is supported by the Austrian Science Fund (FWF-P24551 and I-2805-B29) and partially by the Vienna Graduate School of Population Genetics (FWF-W1225).

## Appendix A The Instantaneous Rate Matrix *Q_revPoMo_*

Let 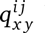 be the rate of a jump from {*ix*, (*N* − *i*)*y*} to {*jx*, (*N* − *j*)*y*}. We can summarize *Q_revPoMo_* as

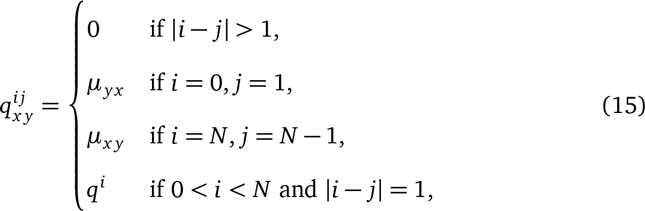

where

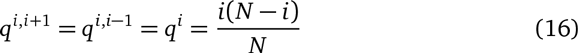

(Section 2.4), *x* ≠ *y* and the diagonal elements (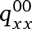 or 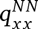 and 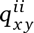, 0 < *i* < *N*) are defined such that the respective row sum is 0.

## Appendix B Derivation of 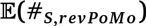

This section derives the expected number of substitutions of revPoMo (Section 2.8). If we denote 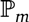 to be the probability of an event to be a mutation, the expected number of substitutions is

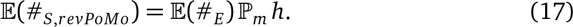

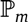 is the ratio of the rate of mutations *r_m_* to the total rate *r* = *r_m_* + *r_s_*. We have

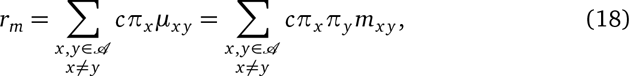

and for the rate of frequency shifts *r_s_*

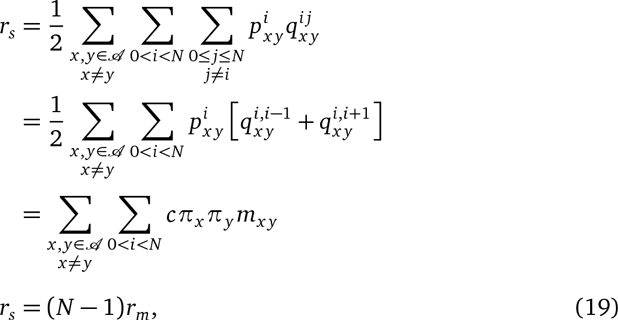

where the 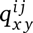 are the rates from {*ix*,(*N* − *i*)*y*} ⟶ {*jx*,(*N* − *j*)*y*} (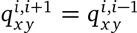; Section S1). Finally,

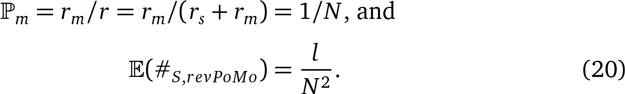

